# Knockout of mitogen-activated protein kinase 3 causes barley root resistance against *Fusarium graminearum*

**DOI:** 10.1101/2022.05.04.490681

**Authors:** Jasim Basheer, Pavol Vadovič, Olga Šamajová, Pavol Melicher, George Komis, Pavel Křenek, Michaela Králová, Tibor Pechan, Miroslav Ovečka, Tomáš Takáč, Jozef Šamaj

## Abstract

The roles of mitogen-activated protein kinases (MAPKs) in plant-fungal pathogenic interactions are less understood in crops. Here, microscopic, phenotyping, proteomic and biochemical analyses revealed that independent TALEN-based knockout lines of *Hordeum vulgare MITOGEN-ACTIVATED PROTEIN KINASE* 3 (*HvMPK3* KO) were resistant against *Fusarium graminearum* infection. When co-cultured with roots of the *HvMPK3* KO lines, *F. graminearum* hyphae were excluded to the extracellular space, the growth pattern of hyphae was considerably deregulated, mycelia development was less efficient and number of appressoria and their penetration potential were significantly reduced. Intracellular penetration of hyphae was preceded by the massive production of reactive oxygen species (ROS) in attacked cells of the wild type, but it was mitigated in the *HvMPK3* KO lines. Suppression of ROS production in these lines coincided with the elevated abundances of catalase and ascorbate peroxidase. Moreover, differential proteomic analysis revealed downregulation of defense-related proteins in wild type, and the upregulation of peroxidases, lipid transfer proteins, and cysteine proteases in *HvMPK3* KO lines after 24h of *F. graminearum* inoculation. Consistently with proteomic analysis, microscopic observations showed an enhanced suberin accumulation in roots of *HvMPK3* KO lines, most likely contributing to the arrested infection by *F. graminearum*. These results suggest that TALEN-based knockout of *HvMPK3* leads to the barley root resistance against Fusarium root rot.

## Introduction

Cereals are a major food staple for the world population, however, there is a substantial reduction in the annual yield because of pathogens, weeds, temperature extremes, high salt concentrations, drought, and arid conditions (Sewelam *et al*., 2016). Diseases caused by fungal pathogens alone are responsible for the destruction of approximately 125 million tons of crops worldwide in the last decade (Vogelgsang *et al*., 2019).

To combat invading fungi, plants recognize microbe-associated molecular patterns (MAMPs), and damage-associated molecular patterns (DAMPs), which are essential for triggering plant immunity (Schwessinger and Zipfel, 2008). MAMPs and DAMPs elicit so-called pattern-triggered immunity (PTI), which is mediated by plant pattern recognition receptors (PRRs) (Huang *et al*., 2016). PTI is characterized by a series of events, including signaling through protein kinase cascades, generation of reactive oxygen species (ROS), calcium ion influx, transcriptional reprogramming, cell wall appositions, and hormonal changes (Kurusu *et al*., 2015). ROS play a dual role during plant immune responses. In addition to signaling functions (Qi *et al*., 2017), their over-accumulation leads to the hypersensitive response, which hinders the progression of the biotrophic pathogens to the plant tissues (Camagna and Takemoto, 2018; Camejo *et al*., 2016). In contrast, necrotrophic pathogens may benefit from nutrients provided by the damaged tissues, and thus ROS accumulation facilitates the invasion of the pathogen into the tissues (Barna *et al*., 2012; Kámán-Tóth *et al*., 2019).

Mitogen-activated protein kinases (MAPKs) are integrated in signaling cascades responsible for conveying signals generated by extracellular and intracellular stimuli (Komis *et al*., 2011). In Arabidopsis and rice, there are several numbers of MAPKs described, of which mostly AtMPK3, AtMPK4, and AtMPK6 and their rice orthologues, are responsible for plant resistance against pathogens (Bigeard *et al*., 2015; Kishi-Kaboshi *et al*., 2010; Meng and Zhang, 2013). The signaling pathway involving AtMPK3/AtMPK6 pair is responsible for the regulation of camalexin biosynthesis (a major phytoalexin found in *Cruciferae* plants) during infection of Arabidopsis by necrotrophic fungal pathogens (Mao *et al*., 2011). However, in the case of bacterial infection, AtMPK4 is responsible for its regulation (Bazin *et al*., 2020). MPK3/MPK6 in different plant species also regulate hypersensitive cell death responses and the generation of reactive oxygen species (Kim and Zhang, 2004; Kroj *et al*., 2003; Liu *et al*., 2007; Ren *et al*., 2002). The importance of MAPK signaling in plant-pathogen interactions is also supported by studies of bacterial effectors, several of which target and inhibit plant MAPK cascades (Cui *et al*., 2010; Zhang and Dong, 2007). Recently, it was shown that knock-out mutations of *HvMPK3* prepared by TALEN technology attenuated the responsivity of barley to bacterial PAMP flagellin 22 (flg22), manifested by decreased abundance of chitinases and pathogenesis-related proteins (Takáč *et al*., 2021).

*Fusarium graminearum* causes a major loss of barley yield due to head blight and root rot diseases. This fungus is considered a hemibiotrophic pathogen, living in the host plants for a short period (hours to several days), before switching to necrotrophic form, which retrieves nutrients from dead cells (Tucker *et al*., 2021, 2019). Although Fusarium head blight was earlier believed as a primary disease caused by *F. graminearum*, the root colonization by this pathogen is currently recognized as very important for immense economic losses. Fusarium root rot causes rapid necrosis, leading to a significant reduction in root growth and biomass, which is accompanied by the progression of the pathogen to the stem base (Smiley *et al*., 2005). The growth-inhibiting impact of the pathogen was assigned to the production of the mycotoxin deoxynivalenol (DON) (Masuda *et al*., 2007). The hyphae colonize intra- and intercellular spaces in the root cortex in sensitive wheat cultivars, while the invasion in resistant cultivar is stopped at the epidermal cells (Wang *et al*., 2015). Barley defense mechanisms against Fusarium root rot are poorly understood. So far, these resistance strategies involved *de novo* biosynthesis of barley root exudates (Lanoue *et al*., 2010), activation of jasmonic acid (JA)- dependent defense genes and genes related to DON detoxification (Wang *et al*., 2018). The role of MAPKs in barley responses to Fusarium has not been elucidated, and their participation in Fusarium-induced signal transduction is unknown.

In the present study, we discovered that independent barley lines with TALEN-based knock-out mutations of *HvMPK3* exhibited higher root resistance to *F. graminearum* and produced less ROS than wild type plants. Fusarium hyphae potential to penetrate the root cells of *HvMPK3* lines significantly decreased. This was accompanied by the attenuation of defense responses in wild type plants and the induction of upregulated levels of cysteine proteases and secretory peroxidases in *HvMPK3* KO lines, as documented by proteomic analysis. Our results indicate that ROS generation, might facilitate the invasion of wild type plants by *F. graminearum*, but elevated abundances of cytosolic ascorbate peroxidase and catalase might reduce ROS levels, thus contributing to the higher resistance of *HvMPK3* KO lines. Finally, the stiffening of the cell walls by suberin deposition likely represents a barrier, which prevents pathogen invasion to the root cells of *HvMPK3* KO lines.

## Results

### Phenotype of wild type and HvMPK3 KO plants infected by Fusarium graminearum

Fungal mycelium developed extensively after infecting the roots of five days old barley seedlings by *F. graminearum* conidia. Ten days after infection, densely developed mycelium surrounded seeds and basal parts of the root system in wild type (WT) plants (Figure S1A). Roots of WT plants were largely arrested in their growth (Figure S1A, black arrows). In contrast, the root system of *HvMPK3* KO-A (Figure S1B), *HvMPK3* KO-B (Figure S1C), and *HvMPK3* KO-D (Figure S1D) lines inoculated by *F. graminearum* developed well without any visible reduction of root growth. In addition, the development of *F. graminearum* mycelia itself was inhibited, particularly around the seeds and roots (Figure S1B-D). Dark brown coloration of *F. graminearum* mycelia appeared close to the seeds and the basal parts of the root system of *HvMPK3* KO lines (Figure S1B-D). On WT genotypes, however, *F. graminearum* mycelia showed pale white color and massive coverage of infected plants (Figure S1A).

Inoculation of control seedling roots (Figure 1A-D) with *F. graminearum* spores led to their germination and subsequent colonization of roots by growing mycelia. In a period of 24 h after inoculation (Figure 1E-H), the root apex of WT plants was massively invaded by GFP-expressing *F. graminearum* hyphae (Figure 1E). However, only sparse hyphae were present on the root surface of *HvMPK3* KO lines (Figure 1F-H). Propidium iodide (PI) staining of cell walls in living cells of uninfected plants (Figure 1A-D) served as a marker of root tissue organization, while accumulation of PI in nuclei in dead cells of infected plants (Figure 1E-H) indicated their mortality after penetration by *F. graminearum* hyphae. The cell death in the epidermis of WT roots colonized by *F. graminearum* was evident (Figure 1E), while the amount of dead epidermal cells in inoculated roots of *HvMPK3* KO lines was considerably low (Figure 1F-H). The quantitative analysis of cell death rate revealed a statistically significant increase in infected WT plants. In contrast, there was no statistical difference in the number of dead root epidermal cells between uninfected plants and plants analyzed 24h after inoculation in *HvMPK3* KO lines (Figure 1I).

**Figure 1.**
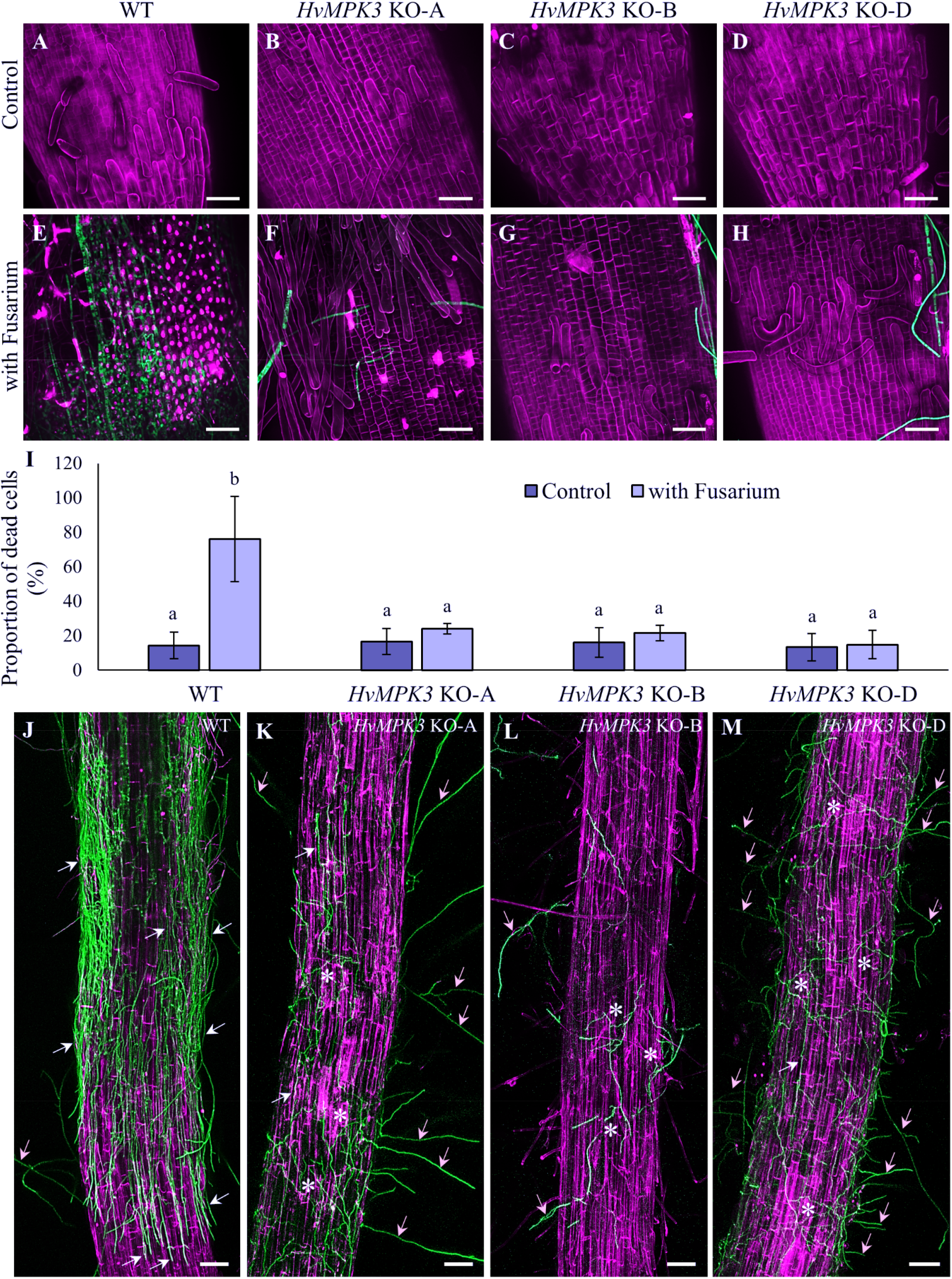
Colonization of root in barley wild type (WT) and *HvMPK3* KO lines by *Fusarium graminearum* mycelia 24h and 48h after inoculation with spores. (**A-H**) Overview of propidium iodide-labeled uninfected root apex of WT (**A**), *HvMPK3* KO-A (**B**), *HvMPK3* KO-B (**C**) and *HvMPK3* KO-D (**D**) lines under control conditions, and 24h after inoculation of roots with spores of GFP-expressing *F. graminearum* in WT (**E**), *HvMPK3* KO-A (**F**), *HvMPK3* KO-B (**G**) and *HvMPK3* KO-D (**H**) lines. Note red propidium iodide staining of cell walls in living cells and nuclei in dead cells. (**I**) Quantitative evaluation of the proportion of dead cells in control and infected root apices among tested barley lines. Data are presented as means ± SD; n = 20 roots/line, 70-90 epidermal cells evaluated in each root, error bars represent standard deviations. Different lowercase letters above the error bars (**I**) represent statistical significance according to one-way ANOVA and subsequent LSD test at *p* value < 0.05. (**J-M**) Roots of WT (**J**), *HvMPK3* KO-A (**K**), *HvMPK3* KO-B (**L**) and *HvMPK3* KO-D (**M**) lines stained with propidium iodide 48h after inoculation with spores of GFP-expressing *F. graminearum*. Hyphae growing on root surface in parallel orientation with elongated root epidermal cells are indicated by white arrows, hyphae growing away and not touching roots are indicated by yellow arrows, zones occupied by malformed mycelia with wavy growth pattern, growing perpendicularly to elongated root epidermal cells are indicated by asterisks. Scale bars: 50 μm (**A**-**H**) and 100 μm (**J**-**M**).

Examination of plants 48h after inoculation with *F. graminearum* showed that roots of WT plants were massively colonized by the fungal mycelium (Figure 1J), but mycelium was much less developed around the infected area on the root surface in *HvMPK3* KO lines (Figure 1K-M). Surprisingly, this fact was related to different growth pattern of hyphae, including considerable changes in their density, but also in the shape of developing mycelium. On the surface of WT roots, the hyphae were growing in parallel with longitudinal root axis and closely associated with the surface of root epidermal cells (Figure 1J). In contrast, hyphae were growing away of the roots and often without touching the root surface of *HvMPK3* KO lines (Figure 1K-M). In addition, mycelium developed around *HvMPK3* KO roots was malformed with unusual wavy growth pattern of hyphae, and showed changed growth direction oriented perpendicularly to the axis of elongated root epidermal cells (Figure 1K-M). Using analysis of the angular distribution of growing mycelia, we quantitatively determined a degree of anisotropy of hyphae distribution at the root surface. In the WT roots, the graph shows almost uniformly longitudinal orientation of hyphae relative to the root longitudinal axis (Figure S2A). Conversely, mycelia around root surfaces of *HvMPK3* KO lines showed more random organization (Figures S2B-S2D). Quantitative evaluation revealed higher average values for angular distribution of hyphae on WT roots (Figure S2E), suggesting a high degree of anisotropy, characterized by hyphae arrangement in one orientation. Opposite, statistically significant lowering of values determining a high degree of isotropy characterized by uniform distribution of hyphae with no prevalent orientation, was found in *HvMPK3* KO lines (Figure S2E). We also analysed fluorescence skewness, defining a pattern of fluorescence intensity distribution on the root surface and comparing how a ratio between high and low fluorescence intensities distribution is changing. This analysis revealed a high degree of fluorescence uniformity (referred by lower values) on the surface of WT roots (Figure S2F), but a high degree of non-uniformity of the fluorescence intensity distribution (characterized by distinct dark and bright fluorescent areas and referred by higher values) on the surface of *HvMPK3* KO roots (Figure S2F).

In comparison to uninfected WT plants (Figure 2A), analysis of root system phenotypes of *F. graminearum*-treated WT plants (10 days after inoculation) revealed a reddish-brown coloration and malformation of some roots (Figure 2E). However, a comparison of roots of uninfected *HvMPK3* KO lines (Figure 2B-D) with infected ones (Figure 2F-H) showed no change. Roots in the basal part of the root systems of *HvMPK3* KO lines did not exhibit a reddish-brown coloration (Figure 2F-H). The morphology (Figure S3) and the length (Figure 2I) of roots measured 10 days after inoculation revealed that *F. graminearum* infection significantly reduced the root length in the WT plants, but there was no significant difference between control *versus* infected *HvMPK3* KO lines (Figure 2I). These data indicate insensitivity of *HvMPK3* KO plants to *F. graminearum* infection in early seedling stages, and sustained root growth without deleterious effects of the pathogen.

**Figure 2.**
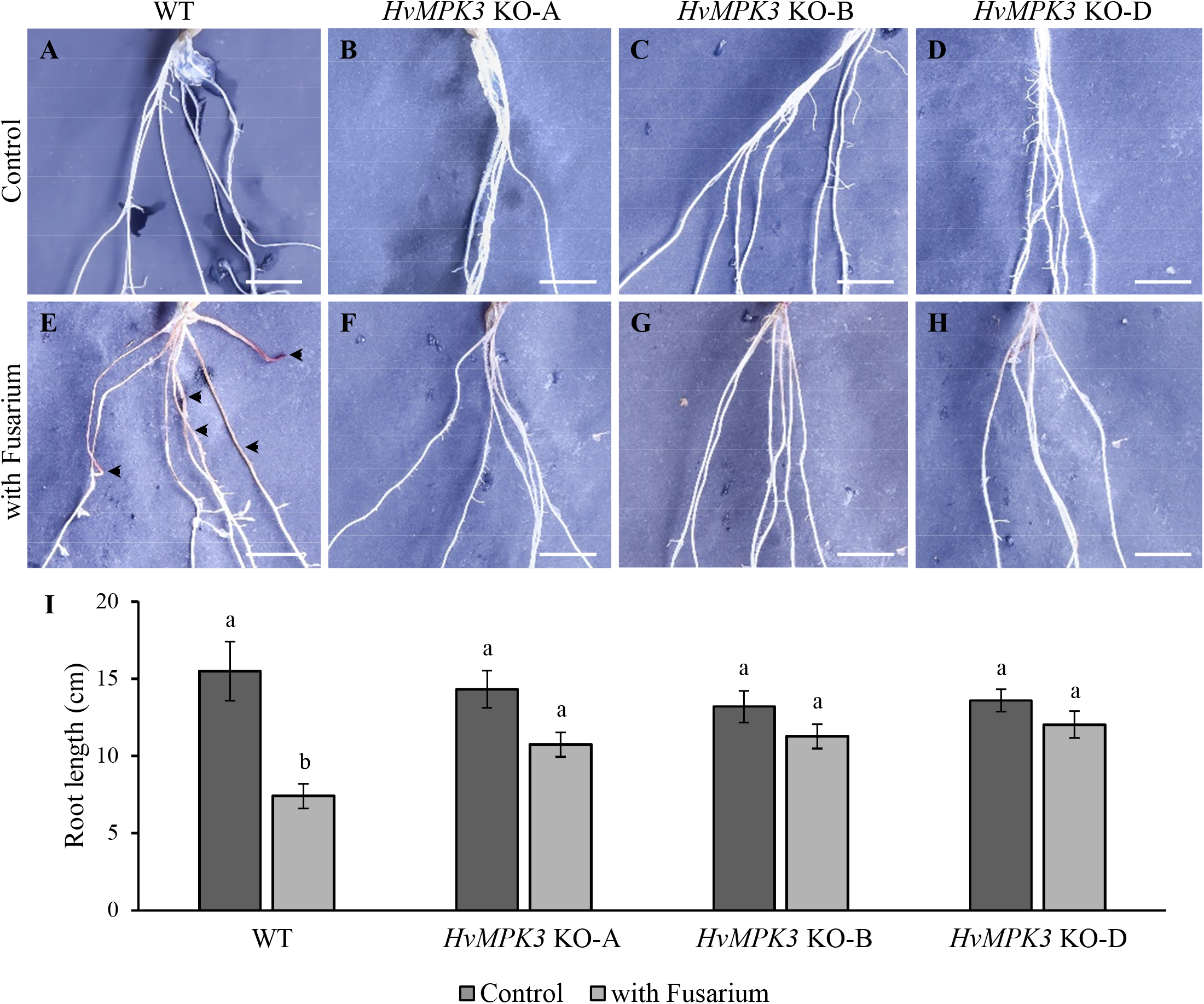
Comparison of root phenotypes in barley wild type (WT) and *HvMPK3* KO lines between non-treated and *Fusarium graminearum-treated* plants 10 days after inoculation with spores. (**A-H**) Morphology of roots in the basal part of the root system of the uninfected plant of WT (**A**), *HvMPK3* KO-A (**B**), *HvMPK3* KO-B (**C**) and *HvMPK3* KO-D (**D**) lines under control conditions, and 10 days after inoculation in WT (**E**), *HvMPK3* KO-A (**F**), *HvMPK3* KO-B (**G**) and *HvMPK3* KO-D (**H**) lines. The extent of infection and the rate of root damage is indicated by a reddish-brown coloration originating from the *F. graminearum* mycelia. Note apparent morphological changes in roots of WT (**E**, arrowheads) that are not present in roots of *HvMPK3* KO-A (**F**), *HvMPK3* KO-B (**G**) and *HvMPK3* KO-D (**H**) lines. Uncropped original images of the tested barley plants are presented in Figure S3. Scale bars: 1 cm. (**I**) Quantitative evaluation of the root length in WT and *HvMPK3* KO lines without infection (Control), and 10 days after infection with *F. graminearum* spores (with Fusarium). Data are presented as means ± SD; n = 30 plants/line, error bars represent standard deviations. Different lowercase letters above the error bars (**I**) represent statistical significance according to one-way ANOVA and subsequent LSD test at *p* value < 0.05.

Growth of the mycelium on the root surface precedes the hyphae invasion to the root cells by cell wall penetration. Volumetric fluorescence visualization of GFP-expressing *F. graminearum* hyphae on the root surface allowed not only qualitatively determine the distribution and density of mycelium, but also its penetration ability. Therefore, we documented the depth of hyphal invasion by color-coded fluorescence intensity distribution along the Z-axis of imaging. In WT roots 48h after inoculation with spores, such analysis revealed formation of appressoria and infection hyphae invading the root epidermal cells. Deeper position of appressoria on cell surface and intracellular location of invading hyphae were clearly distinguished from root surface mycelium according to distinct fluorescence colorcoding (Figure 3A, B). In sharp contrast, the same analysis did not reveal penetration and intracellular infection hyphae formation in the root epidermis of *HvMPK3* KO lines (Figure 3C-H). Overall microscopic analysis revealed formation of dense mycelium on the surface of WT roots, with frequent infection hyphae penetration to the root epidermal cells (Figure S4A, B). In contrary, low-density surface mycelium with different growth pattern, and no hyphae penetration in the root epidermal cells were revealed in *HvMPK3* KO lines (Figure S4C-H). Moreover, occasional *F. graminearum* appressoria developed on root surfaces of *HvMPK3* KO lines were often abnormal in shape and size (yellow arrows in Figure 3C-H). Differences in appressoria morphology between *F. graminearum* mycelium growing on the surface of WT roots (Figure 3A, B, 4A) and roots of *HvMPK3* KO lines (Figure 3C-H, 4B-D) prompted us to analyze appressoria formation capacity, which may directly influence also their penetration ability to the root epidermal cells. We have found significantly lower numbers of appressoria (Figure 4E) and penetrating hyphae (Figure 4F) on the surface of *HvMPK3* KO roots compared to the WT. Therefore, data from phenotypical analysis revealed different responses of WT and *HvMPK3* KO roots to *F. graminearum* infection.

**Figure 3.**
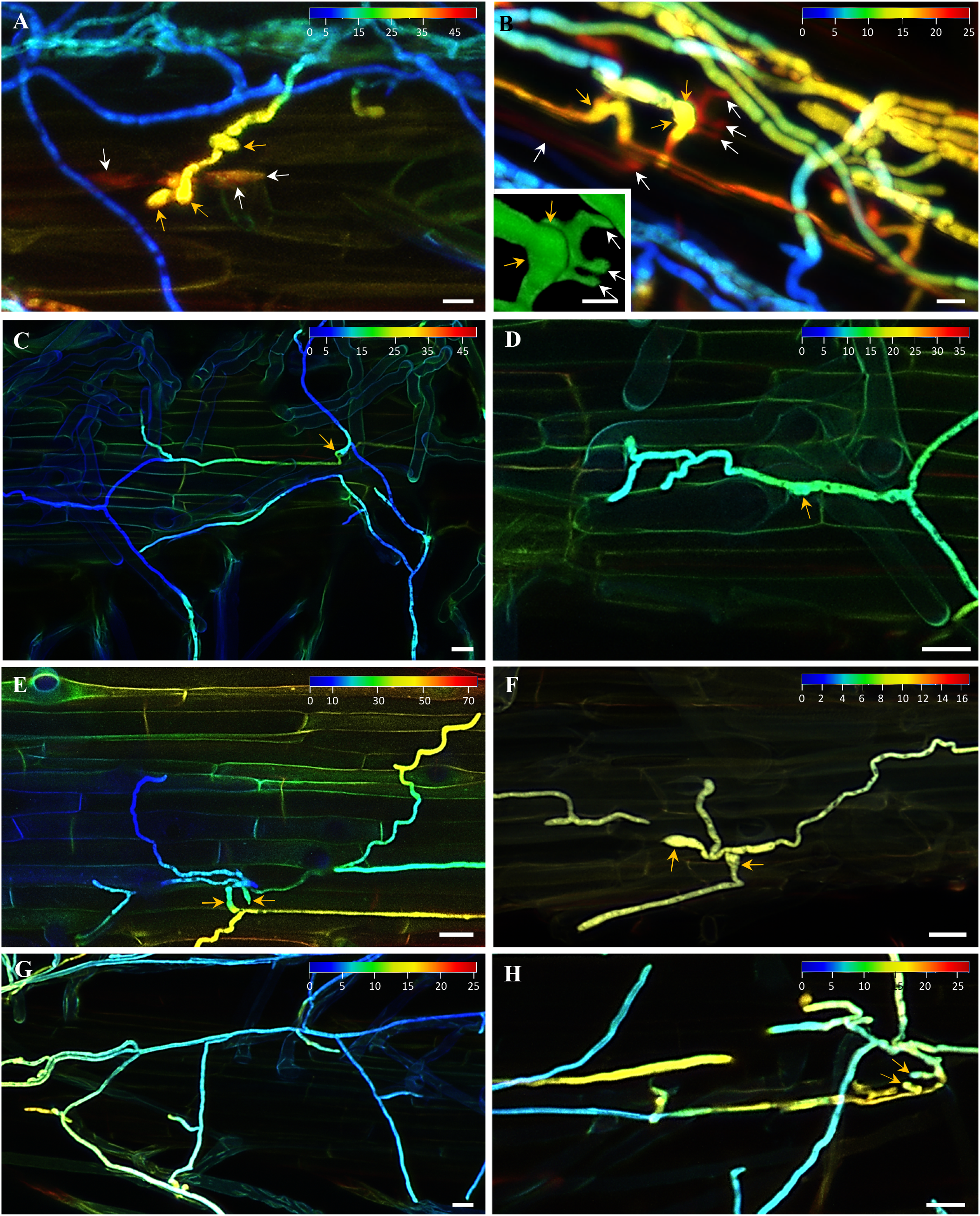
Visualization of volumetric fluorescence signal distribution of *F. graminearum* hyphae growing on root surface of barley wild type (WT) and *HvMPK3* KO lines 48h after inoculation with spores. Depth of the hyphae localization is indicated by a color-coding depicted on the root surface by blue color and deep inside the root tissue by red color. The color range in each image is indicated in μm. (**A, B**) Formation of appressoria (yellow arrows) and infection hyphae (white arrows) penetrating cell wall of epidermal cells in roots of WT plants. Inset in (**B**) shows detailed 3D visualization of appressorium (yellow arrows) and infection hyphae (white arrows). (**C-H**) Developed mycelia on root surface of *HvMPK3* KO-A (**C, D**), *HvMPK3* KO-B (**E, F**) and *HvMPK3* KO-D (**G, H**) lines. Appressoria are indicated by yellow arrows, cell wall penetration and formation of infection hyphae were not detected by color-coded analysis. Scale bars: 10 μm (**A**, **B**) and 20 μm (**C**-**H**).

**Figure 4.**
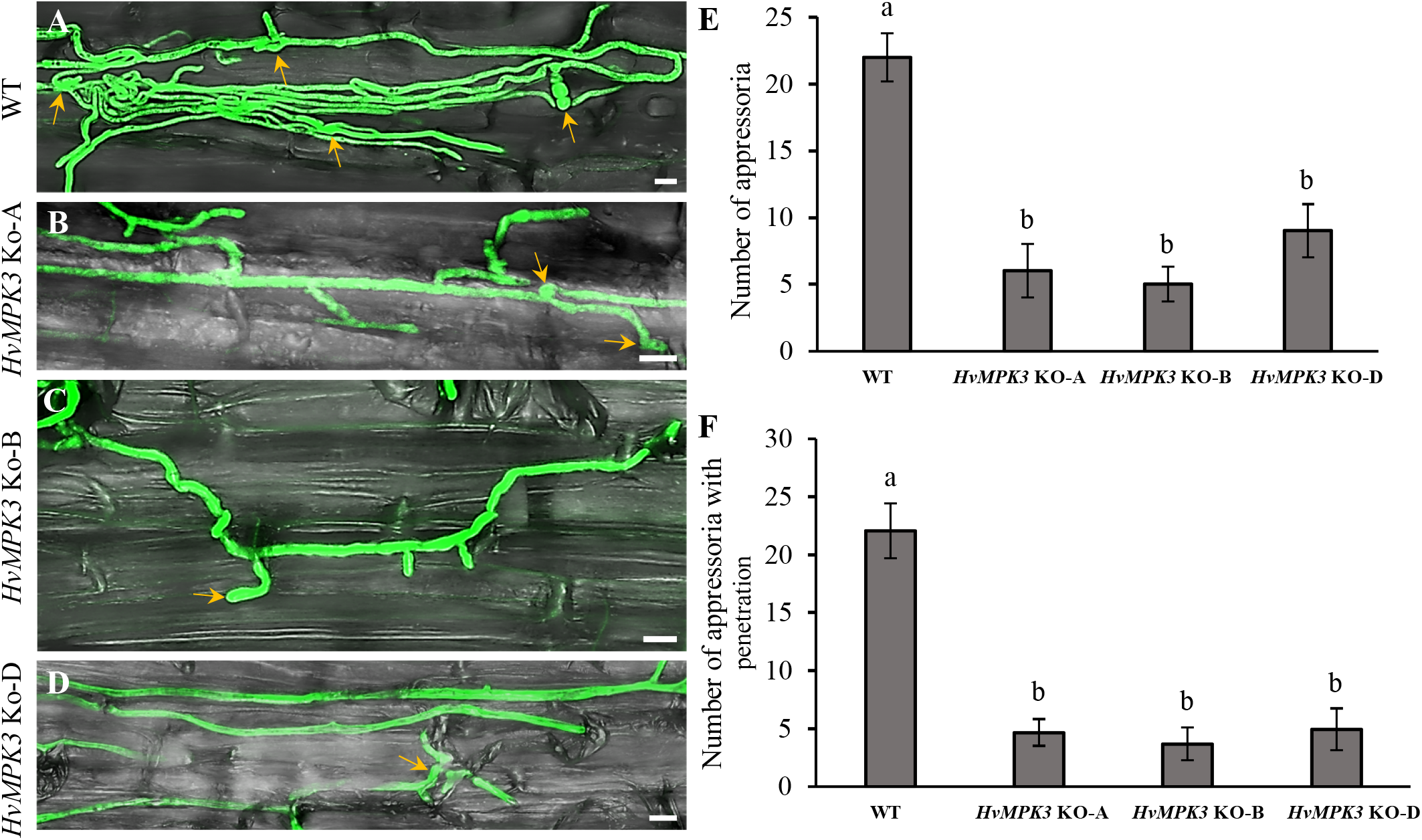
Morphology of *F. graminearum* appressoria, their number and the mycelium penetration potential on root surface of barley wild type (WT) and *HvMPK3* KO lines 48h after inoculation with spores. **(A-D)** Mycelium density and the appressoria morphology 48h after inoculation with *F. graminearum* spores on the root surface of WT **(A)**, *HvMPK3* KO-A (**B**), HvMPK3 KO-B (**C**) and HvMPK3 KO-D (**D**) lines. Appressoria are indicated by yellow arrows. Scale bars: 20μm. **(E)** Quantitative evaluation of the average number of appressoria developed on the surface of infected roots. **(F)** Quantitative evaluation of the average number of the penetration events on the surface of infected roots. Data are represented as mean ± SD; n = 10 images/line, error bars represent standard deviations. Different lowercase letters above the error bars represent statistical significance according to one-way ANOVA and subsequent LSD test at *p* value < 0.05.

### Fusarium-induced ROS production in barley roots

ROS production in infected plant cells is associated with the early recognition of pathogen at the apoplast, and is accompanied by sudden oxidative burst (Qi *et al*., 2017). Therefore, ROS production in roots of WT and *HvMPK3* KO lines was analysed by CM-H_2_DCFDA fluorescence detection 24h after inoculation with *F. graminearum* spores. This analysis revealed a substantially higher ROS accumulation in WT (Figure 5A), compared to the *HvMPK3* KO lines (Figure 5B-D). Consequently, quantitative evaluation of the fluorescent staining levels showed that ROS accumulation was substantially higher in the infected WT roots than in infected roots of *HvMPK3 KO* lines (Figure 5E). Independent spectrophotometric examination of ROS levels in the infected WT and *HvMPK3* KO lines showed results consistent with these microscopic observations (Figure S5). In WT plants, ROS production in cells attacked by fungus preceded fungal hyphae penetration (Figure 5F). Such cells in the preinvasion stage were still alive and accumulated a high amount of ROS, while other surrounding cells with absent ROS fluorescence signal were dead (Figure 5F). In the root apex of WT plants, ROS production and accumulation occurred in cells not yet penetrated with *F. graminearum* hyphae, while in most of the root epidermal cells ROS production was not detected (Figure 5G). Based on the staining of nuclei by PI, these cells were already dead and penetrated by *F. graminearum* hyphae (Figure 5G). Conversely, only a limited number of root epidermal cells appeared dead after PI staining of *HvMPK3* KO lines 24h after inoculation with *F. graminearum* spores (Figure 5H-J). Fluorescence signal for ROS detection was not observed, which was related to the low number of *F. graminearum* hyphae growing around the roots and their minimal contact with the root epidermal cells of *HvMPK3* KO lines (Figure 5H-J).

**Figure 5.**
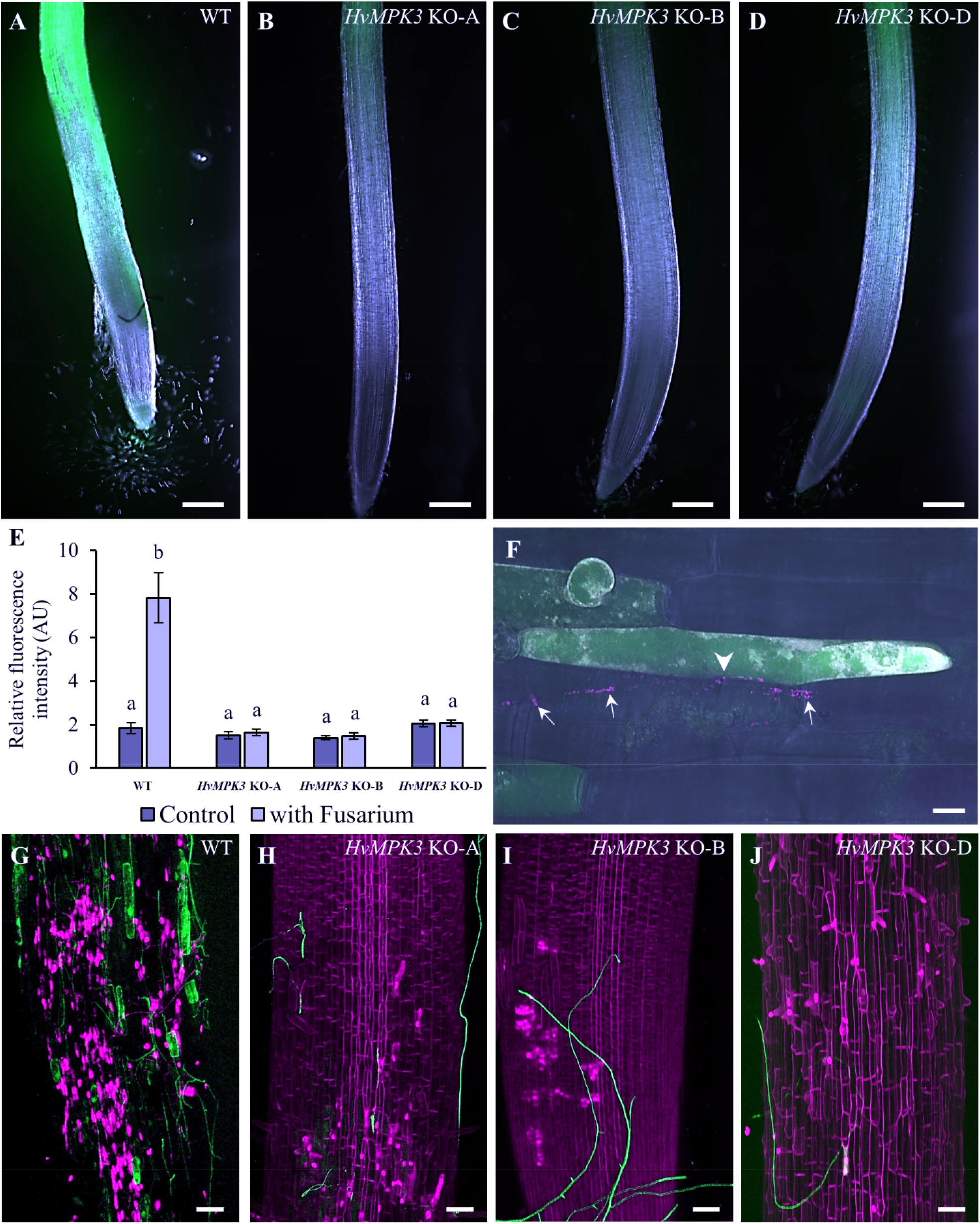
Fluorescence detection of ROS production in roots of barley wild type (WT) and *HvMPK3* KO lines 24h after inoculation with *Fusarium graminearum* spores. (**A-D**) Overview of the root apex labeled by CM-H_2_DCFDA probe 24h after inoculation in WT (**A**), *HvMPK3* KO-A (**B**), *HvMPK3* KO-B (**C**) and *HvMPK3* KO-D (**D**) lines. (**E**) Quantitative evaluation of the fluorescence intensity after CM-H_2_DCFDA labeling of control and inoculated roots. Quantification was performed 24h after inoculation with *F. graminearum*. Data are presented as means ± SD; n = 15 plants/line, error bars represent standard deviations. Different lowercase letters above the error bars represent statistical significance according to one-way ANOVA and subsequent LSD test at *p* value < 0.05. (**F**) Increased ROS production in root epidermal cells of WT plant treated with mCherry-expressing *F. graminearum* (white arrows) for 24h. Note the close contact of ROS-producing cell and the mycelia (white arrowhead) before infection. (**G-J**) Local view of the infected root zone 24h after inoculation with GFP-expressing *F. graminearum* and subsequently labeled by CM-H_2_DCFDA for ROS and by propidium iodide for cell walls in living cells and for nuclei in dead cells in WT (**G**), *HvMPK3* KO-A (**H**), *HvMPK3* KO-B (**I**) and *HvMPK3* KO-D (**J**) lines. Scale bars: 1 mm (**A-D**), 50 μm (**G-J**), 10 μm (**F**).

### Differential proteomic analysis

To elucidate resistance mechanisms of *HvMPK3* KO lines against *F. graminearum* at molecular level, root proteomes of WT and *HvMPK3* KO lines infected with *F. graminearum* were compared to mock controls 24h after infection.

In total, we have identified 133 differentially abundant proteins in *F. graminearum-* treated WT plants, of which 43 were upregulated and 90 were downregulated in infected *versus* control plants. In *HvMPK3* KO lines, 94 differentially abundant proteins were found, from which 47 were upregulated and the same number was downregulated in infected *versus* control plants (Tables S1 and S2). We evaluated the differential proteomes of WT and *HvMPK3* KO lines using GO annotation analysis (Figure S6-S7).

Here we compared the previously published differential proteome of *HvMPK3-KO* lines exposed to flg22 (Takáč *et al*., 2021) with the differential proteome of the same lines treated with Fusarium. In contrast to flg22, Fusarium caused the differential abundance of lipid transfer proteins (LTPs), histone isoforms and proteins involved in reactive oxygen species regulation (Figure 6). In more detail, abundances of histone isoforms were upregulated in WT, but downregulated in *HvMPK3* KO lines treated with Fusarium (Figure 6). LTPs, a protein family affected by Fusarium but not by flg22, were distinctly affected in KO *HvMPK3* lines compared to WT (Figure 6). Putative lipid transfer-like protein DIR1 and non-specific LTP1 were upregulated while YLS3 was downregulated in infected WT lines. In *HvMPK3* KO lines, Fusarium caused upregulation of two VAS lipid transfer-like proteins and downregulation of CW21 non-specific LTP.

**Figure 6.**
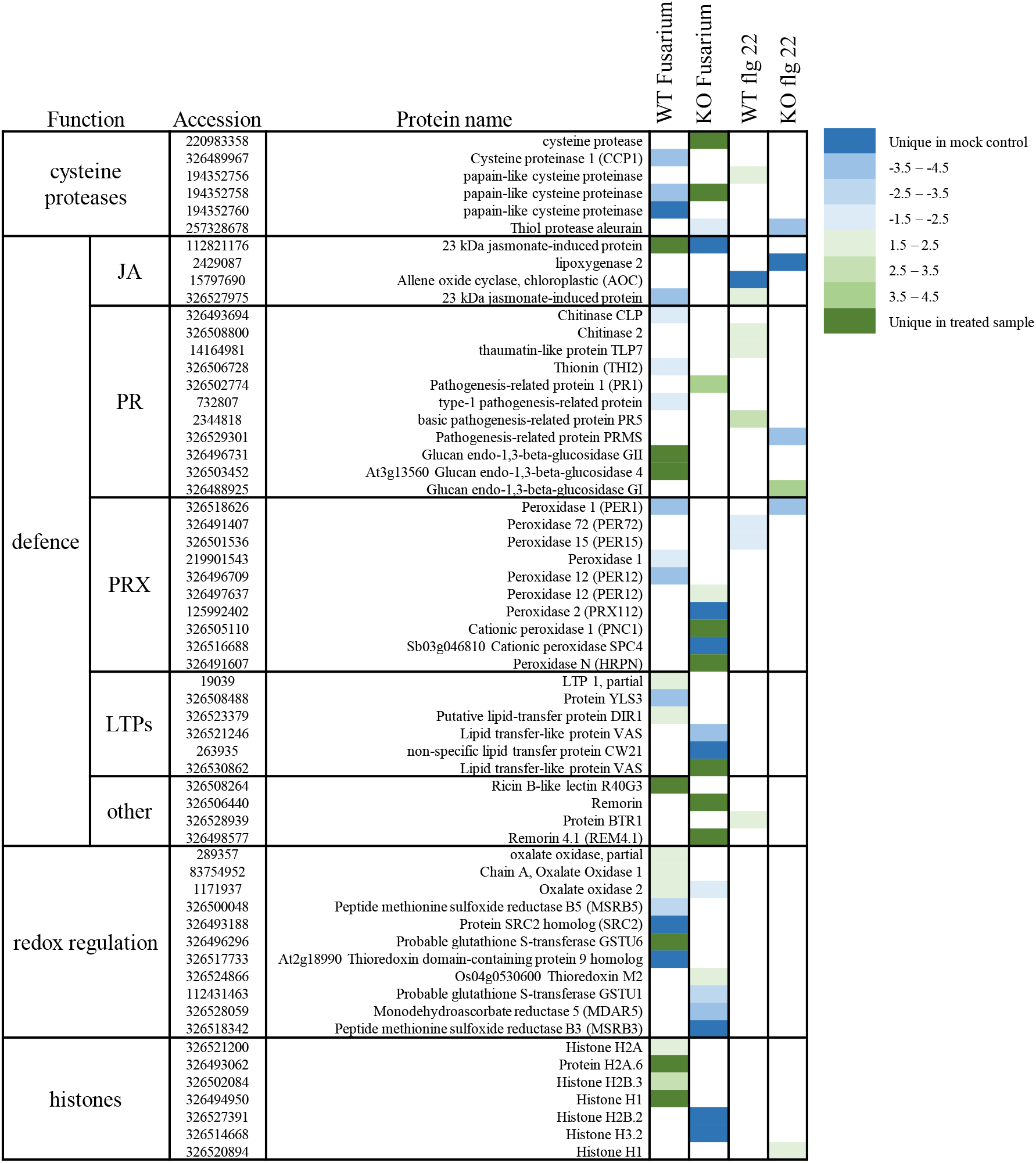
Heat map depicting the fold changes of defence-related proteins, histones, and proteins involved in redox regulation differentially regulated in WT and *HvMPK3* KO plants in response to *Fusarium graminearum* (this study) and flagellin 22 (flg22) treatment (Takáč *et al*., 2021). Fold changes are presented in color as indicated by the color key on the right. Proteins unique for either control or treated samples were found solely in the proteomes of control or treated samples, considering all analyzed replicates. JA = jasmonic acid; PR = pathogenesis related proteins; PRX = peroxidases; LTPs = lipid transfer proteins.

Concerning proteins involved in redox regulation, peptide methionine sulfoxide reductase B5, thioredoxin domain-containing protein 9 and glutathione S-transferase GSTU6 were downregulated in WT (Figure 7). Peptide methionine sulfoxide reductase B3 and glutathione S-transferase GSTU1 were negatively affected by Fusarium treatment in *HvMPK3* KO lines. Monodehydroascorbate reductase 5 (involved in ascorbate regeneration) was downregulated, while thioredoxin M2 (a chloroplastic redox buffering protein) was upregulated in *HvMPK3* KO plants (Figure 7).

**Figure 7.**
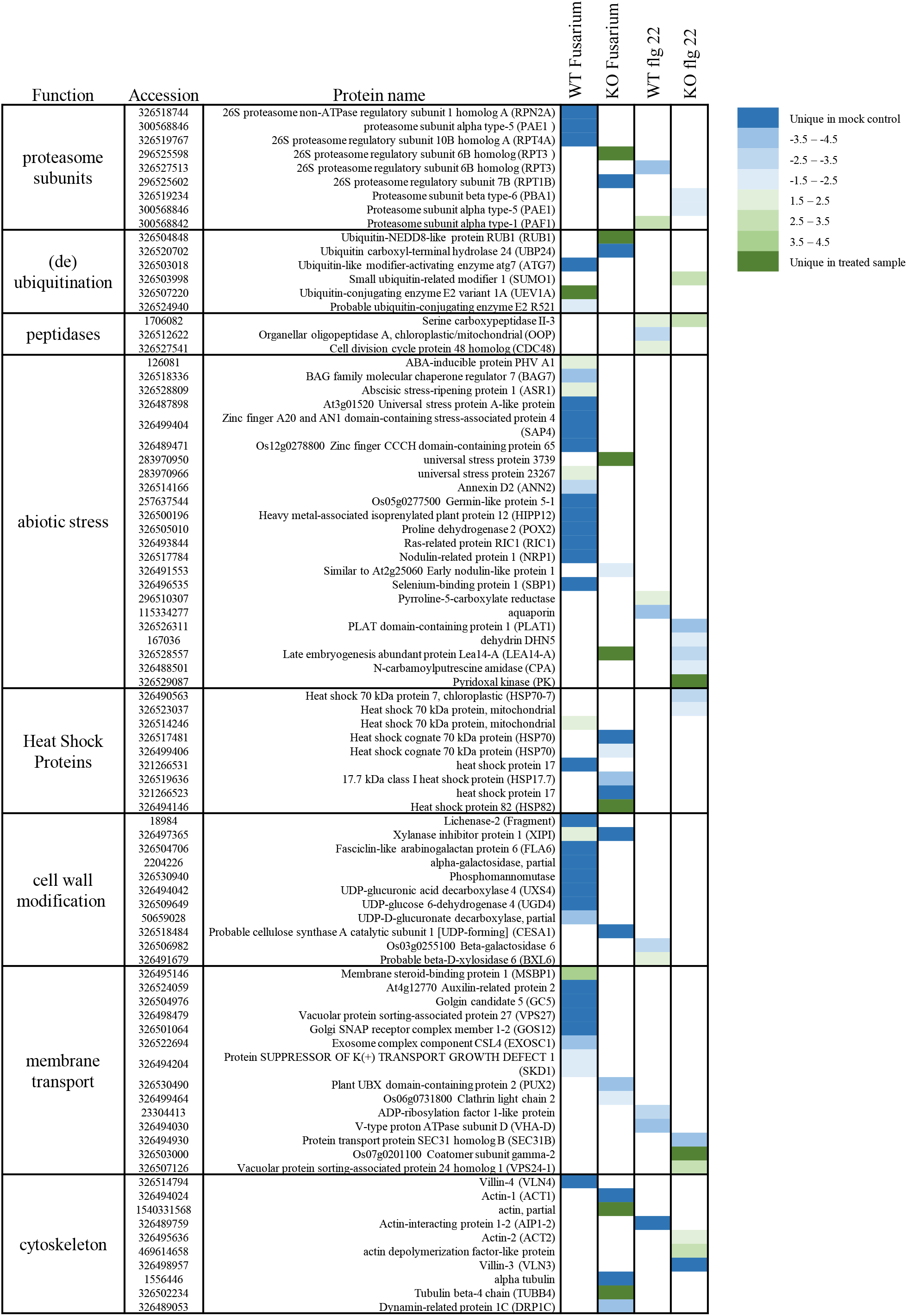
Heat map depicting the fold changes of protein involved in protein degradation, abiotic stress response, cell wall modification, membrane transport and cytoskeleton, differentially regulated in WT and *HvMPK3* KO plants in response to *Fusarium graminearum* (this study) and flagellin 22 (flg22) treatment (Takáč *et al*., 2021). Fold changes are presented in color as indicated by the color key on the right. Proteins unique for either control or treated samples were found solely in the proteomes of control or treated samples, considering all analyzed replicates.

Fusarium also deregulated secretory peroxidases in barley. Proteome-wide, *HvMPK3* KO lines contained five secretory peroxidases and WT only three. Notably, abundances of secretory peroxidases were decreased in WT, while three out of five secretory peroxidases were upregulated in the *HvMPK3* KO lines. Using specific activity staining on native gels, we have found that *HvMPK3* KO lines exhibited also increased activities of three peroxidase isoforms (Rf 0.63, 0.90, 0.98), as compared to the WT (Figure 8A, B). To assess the abundance of major H_2_O_2_-decomposing enzymes, L-ascorbate peroxidase and catalase were analyzed using immunoblot assays. We detected cytosolic APX (cAPX) in the barley root extracts employing anti-ascorbate peroxidase (APX) antibody (Figure 8C, D). Remarkably, the abundance of cAPX increased in *HvMPK3* KO lines, while it decreased in WT plants. In addition, we observed an increase of catalase abundance in both analyzed *HvMPK3* KO lines, while it slightly decreased in WT (Figure 8E, F). These results suggest that *HvMPK3* KO lines likely exert an increased capacity to decompose H_2_O_2_ during *F. graminearum* infection.

**Figure 8.**
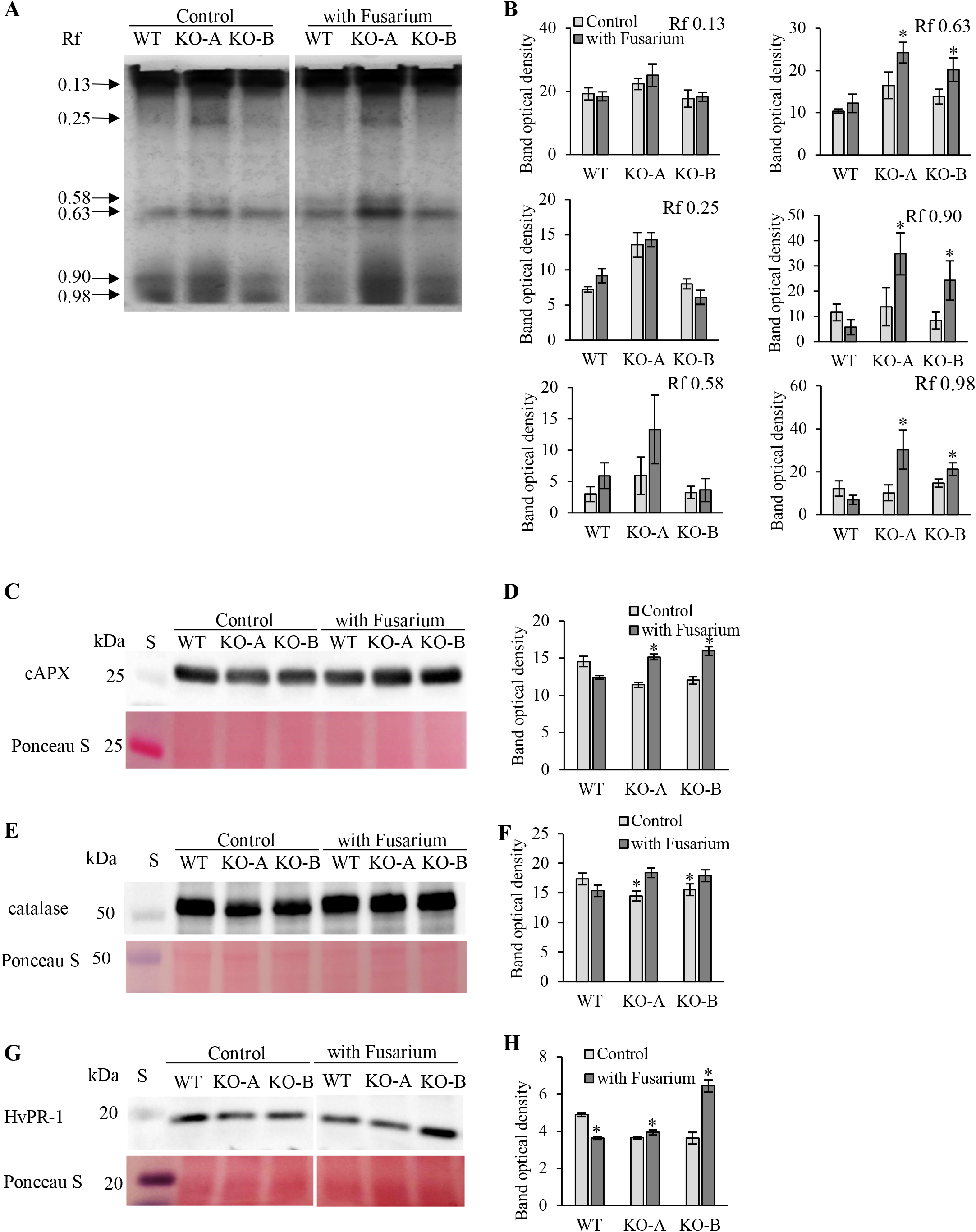
Specific activity of peroxidases and abundances of ascorbate peroxidase and catalase in roots of barley wild type (WT) and *HvMPK3* KO lines (KO-A and KO-B) 24h after inoculation with *Fusarium graminearum*. (**A**) Images of native polyacrylamide gels showing activity of peroxidases. (**B**) Graphs showing quantifications of band optical intensities in (A). (**C, E, G**) Immunoblotting analyses of cytosolic ascorbate peroxidase (cAPX; C), catalase (E) and pathogenesis related protein 1 (HvPR-1; G). (**D, F, H**) Graphs showing quantifications of band optical intensities in (C, E and G). Data are presented as means ± SD; error bars represent standard deviations. Asterisks above the error bars represent statistical significant differences between control and treated samples according to Student’s t-test at p value < 0.05. Uncropped, full original images of the gels are documented in Figure S8.

Moreover, cysteine proteases were more affected by Fusarium as by flg22. They were downregulated in WT but mostly upregulated in *HvMPK3* KO lines. Next we identified more proteins belonging to the family of heat shock proteins in the Fusarium-induced differential proteome. The majority of them was downregulated in *HvMPK3* KO lines.

Proteins involved in cell wall modifications and abiotic stress response were more and preferentially affected by Fusarium as compared to flg22, and these protein groups mostly showed reduced abundances in WT. Finally, enzymes with ubiquitination and deubiquitination activities were also affected by Fusarium, but without differences between WT and *KO HvMPK3* lines.

As *HvMPK3* KO lines exhibited reduced colonization of roots by *F. graminearum* hyphae, we focused on proteins predicted to be localized to extracellular space (Table S3). These included three cysteine proteinases, which were upregulated in *HvMPK3* KO lines, and downregulated in WT. Furthermore, PR proteins (PR-1, thionin 2, glucan endo-1,3-beta-glucosidase 4) were found downregulated in WT. Notably, PR-1 protein was upregulated in *HvMPK3* KO lines. Immunoblotting analysis validated this differential abundance of PR-1 in WT and *HvMPK3* KO lines (Figure 8G, H). Some of the above-mentioned LTPs were predicted as extracellular, including DIR1 putative lipid transfer-like protein, non-specific LTP1 and two VAS lipid transfer-like proteins (Table S3). LTPs together with another extracellular protein, GDSL esterase/lipase At5g33370, are involved in cutin and suberin formation in plants (Edqvist *et al*., 2018; Hong *et al*., 2017). Therefore, we examined suberin accumulation in roots of *F. graminearum*-treated WT and *HvMPK3* KO lines using specific fluorescent dye fluorol yellow 088. This histochemical analysis showed stronger suberin deposition to the root surface cell layers of non-infected *HvMPK3* KO lines (Figure 9B, C, G), but not in WT plants (Figure 9A, G). The suberin histochemical staining at root surface cell layers was considerably enhanced only in *HvMPK3* KO lines after *F. graminearum* infection (Figure 9D, E, F, G). In addition, the suberin staining also increased in the root endodermis and stele of *HvMPK3* KO lines (Figure 9B, C, E, F). In contrast, histochemical staining of lignin with basic fuchsine showed an opposite tendency, and no differences in lignin staining were observed under control conditions (Figure 10A-C, G). Nevertheless, treatment with *F. graminearum* caused over 54,5% decrease of lignin staining in WT roots, but only 18,5-28% decrease in *HvMPK3* KO lines (Figure 10D-G). These results suggest that a suberized cell wall barrier against *F. graminearum* invasion was built up at the surface layers of *HvMPK3* KO roots.

**Figure 9.**
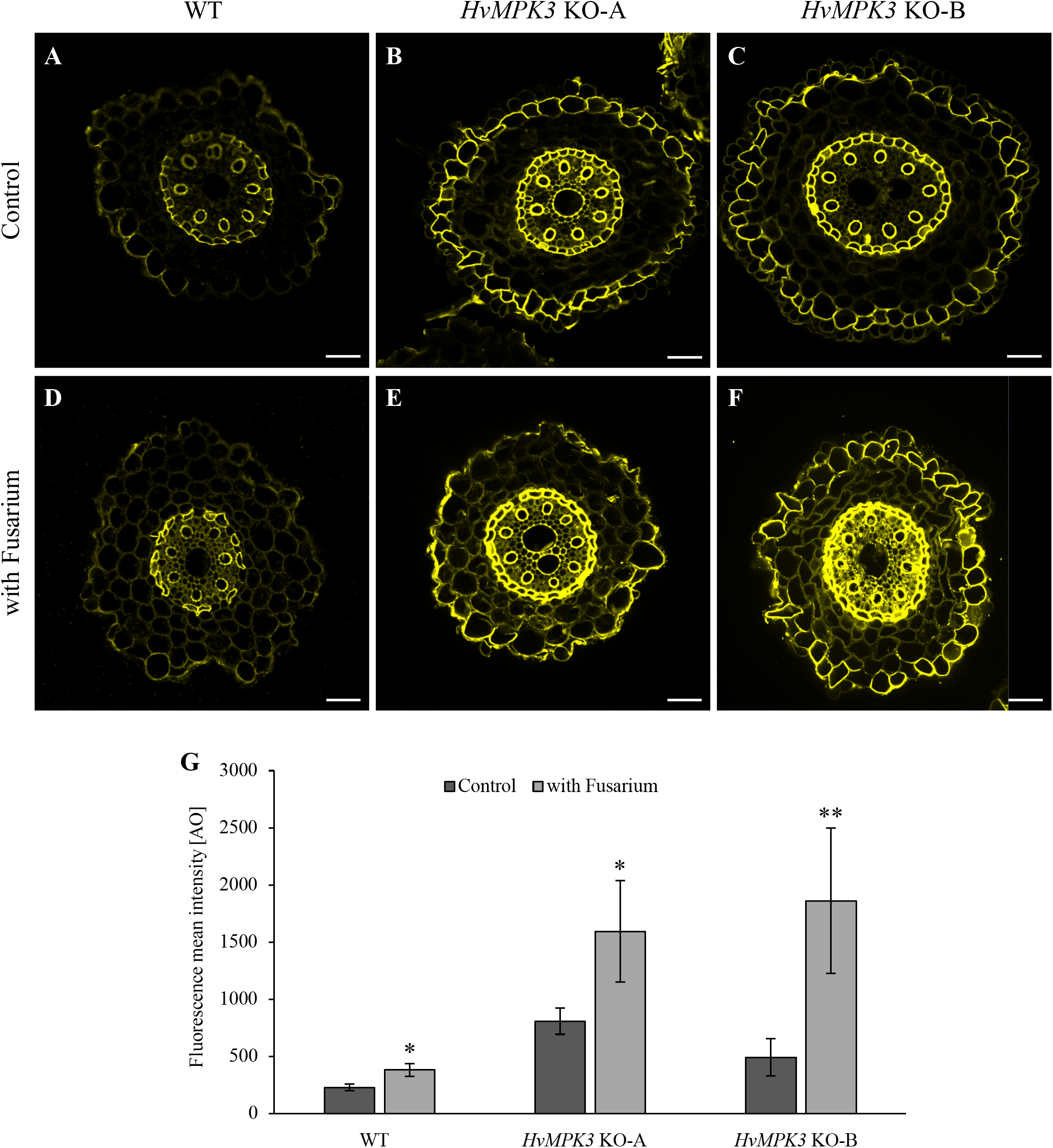
Suberin staining of barley roots 24h after inoculation with *Fusarium graminearum* spores. Freehand cross-sections of roots were stained with fluorescent suberin dye Fluorol Yellow 088 under control conditions (upper panel) and 24h after inoculation with *F. graminearum* spores (lower panel). (**A, B**) wild type, (**C, D**) *HvMPK3* KO-A line and (**E, F**) *HvMPK3* KO-B line. Note strongly increased suberin staining in surface cell layers, endodermis and xylem of both *HvMPK3* KO lines. (**G**) Quantification of suberin staining in freehand barely root cross-sections. Asterisks indicate statistically significant differences between control and treated samples of appropriate line, * p<0.05, ** p<0.01, Student’s t-test. Scale bars: 50 μm.

**Figure 10.**
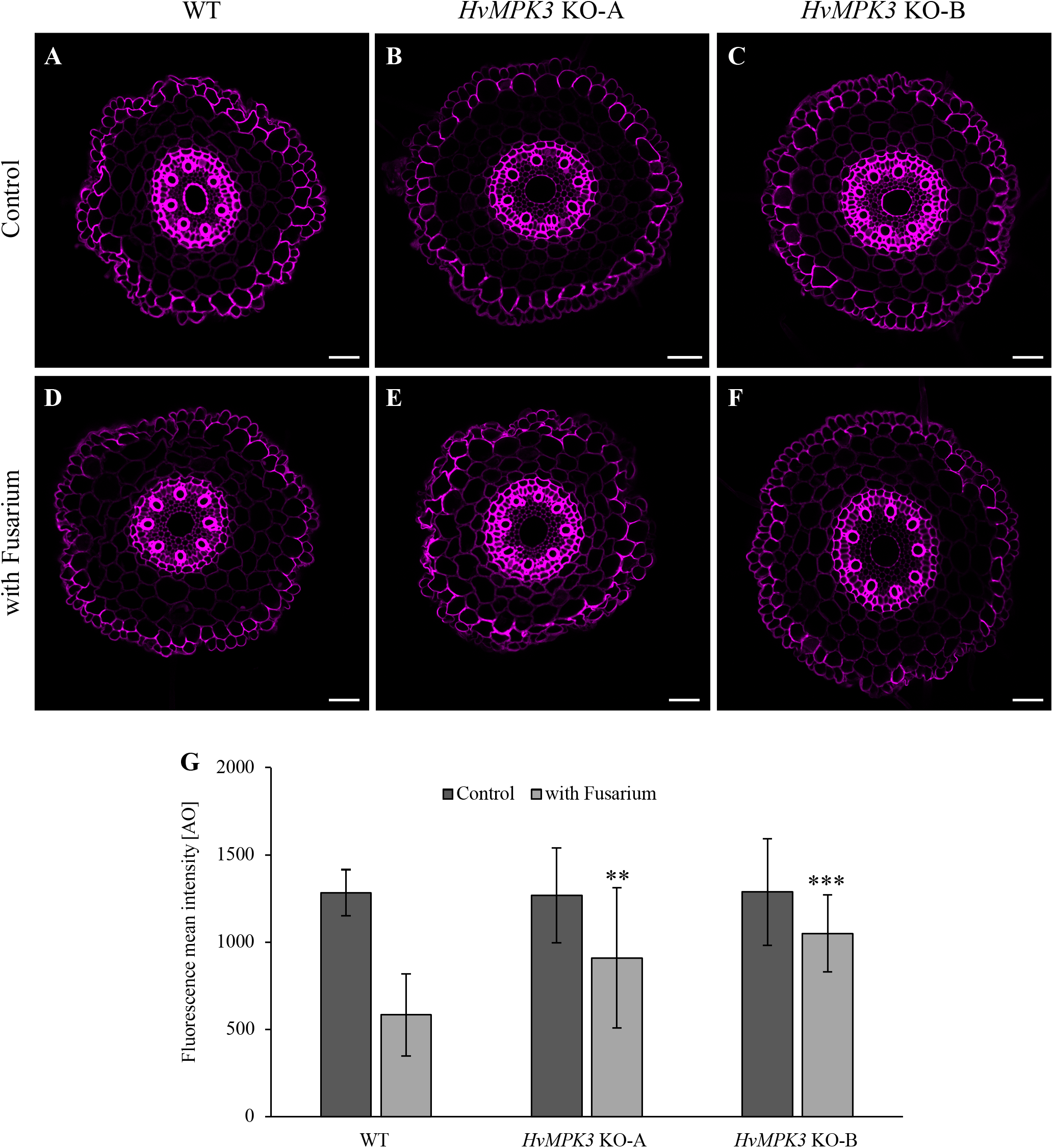
Lignin staining of barley roots 24h after inoculation with *Fusarium graminearum* spores. Freehand cross-sections of roots were stained with fluorescent lignin dye Basic Fuchsine under control conditions (upper panel) and 24h after inoculation with *F. graminearum* spores (lower panel). **(A, B)** wild type, **(C, D)** *HvMPK3* KO-A line and **(E, F)** *HvMPK3* KO-B line. Note strongly increased suberin staining in surface cell layers, endodermis and xylem of both *HvMPK3* KO lines. **(G)** Quantification of suberin staining in freehand barely root cross-sections. Asterisks indicate statistically significant differences between control and treated samples of appropriate line, ** p<0.01, *** p<0.001, Student’s t-test. Scale bars: 50 μm.

In conclusion, proteomic analysis showed that WT exhibited reduced abundances of PR proteins, while the resistance of *HvMPK3* KO lines to *F. graminearum* was accompanied by upregulation of PR proteins, peroxidases, LTPs, cysteine proteinases, proteins involved in suberin formation and it was corroborated by overabundances of cAPX and catalase.

## Discussion

The diversified family of MAPKs lies in the core of plant defense mechanisms due to their significant role in extracellular stimuli perception. In addition, MAPK cascades are responsible for plant resistance to abiotic stresses and exert multiple developmental functions (Bigeard *et al*., 2015; Komis *et al*., 2018). Despite a significant number of MAPK-related studies in dicots like Arabidopsis, the current understanding of MAPK roles in defense of cereal crops against pathogens is still limited. In cereals, MAPKs play either positive or negative roles in immune reactions against fungal pathogens (Křenek *et al*., 2015; Chen *et al*., 2021). In rice, the OsMKK4–OsMPK3/OsMPK6 module participates in transduction of a fungal chitin elicitor signal and regulates defense responses leading to the biosynthesis of diterpenoid phytoalexins and lignin, and progression of plant cell death (Kishi-Kaboshi *et al*., 2010). OsMPK3 negatively regulates the expression of defense genes and the defense reactions against hemibiotrophic blast fungus *Magnaporte grisea* (Xiong and Yang, 2003). It was reported that this negative regulation is mediated through interaction with, and by phosphorylation of OsMPK3 via calcium-dependent protein kinase OsCPK18 (Xie *et al*., 2014). Wheat TaMPK3, unlike TaMPK6, is specifically activated, and its transcript levels as well as abundance are increased during compatible interactions with necrotrophic fungus *Mycosphaerella graminicola* (Rudd *et al*., 2008). This increase is directly correlated with the onset of programmed cell death (PCD) and the generation of related PCD symptoms (Rudd *et al*., 2008). Barley *MPK3* is upregulated in response to the rust fungus *Puccinia hordei* inoculation, especially during effector-triggered immunity (Křenek *et al*., 2015). Remarkably, *HvMPK3* KO lines showed attenuated response to flg22 treatment in terms of defense-related genes such as chitinases, indicating some positive role of HvMPK3 in PTI (Takáč *et al*., 2021).

Although Fusarium root rot is capable to cause severe yield losses, the plant defense mechanism against this disease has not been sufficiently investigated and understood yet. Role of MAPKs in cereal responses to Fusarium were not studied before. The resistance against Fusarium root rot was previously shown to be mediated by hindering the penetration of the fungi at the wheat epidermal cells (Wang *et al*., 2015). Our results suggest that the exclusion of *F. graminearum* to the extracellular space in the roots of *HvMPK3* KO lines is likely caused by suberin deposition to the root surface of these lines, which is also supported by upregulation of proteins involved in suberin formation. We also observed decrease in lignin deposition, similarly as previously published for *F. oxysporum* treatment of flax and *F. solani* treatment of soybean (Lozovaya *et al*., 2006; Wojtasik *et al*., 2016).

When compared to flg22 treatment (Takáč *et al*., 2021), *F. graminearum* infection had more obvious impact on the barley proteome, especially concerning changed abundance of extracellular proteins. This might be assigned to the hemibiotrophic pathosystem of the fungus, eliciting more complex defense mechanisms compared to bacterial elicitor flg22. Hence, this proteomic analysis allowed to gain a more detailed insight into defense mechanisms occurring in *HvMPK3* KO lines and suggested that WT roots were compromised in defense response, most likely due to fungus-secreted toxin effectors. Is was illustrated by the downregulation of PR17c precursor and several isoforms of cysteine proteases, representing well-known pathogen effector targets (Zhang *et al*., 2012, Mueller *et al*., 2013). Cysteine proteases are major regulators of plant defense responses (Misas-Villamil *et al*., 2016) and in contrast to WT, their abundance considerably increased in *HvMPK3* KO lines, indicating that they might mediate resistance of these lines against *F. graminearum*.

Specialized infection structures called appressoria are necessary for the *F. graminearum* infection. It was reported on plants like wheat, rice, corn or barley attacked by different fungal pathogens, including rusts, powdery mildew and blast diseases (Boenisch and Schäfer 2011; Qiu *et al*., 2019). To facilitate host penetration, appressoria of *Magnaporthe oryzae*, generate enormous pressure up to 8.0 MPa that drive physically rupture of the host cell wall (Talbot, 2019). *F. graminearum* can produce multicellular appressoria called infection cushions (Boenisch and Schäfer 2011), whereas the biology of these structures is not completely known. Here, the number of formed and invading appressoria were significantly higher in the case of WT as compared to the *HvMPK3* KO lines. Twenty-four hours after infection, fungal hyphae successfully invaded the epidermal root cells of WT plants, which was preceded by the dramatic ROS increase and accumulation, leading to hyphae penetration to cells. This is entirely consistent with the previously described model of hemibiotrophic and necrotrophic fungal pathosystems (Desmond *et al*., 2008). In contrast, the production of ROS was substantially lower in infected *HvMPK3* KO plants, partly due to prevention of the interaction between the pathogen and the root surface of transgenic plants, and partly also due to their elevated antioxidant capacity mediated by cytosolic ascorbate peroxidase and catalase.

The ROS generated and released during infection can affect both counterparts, the host and the microbe (Desmond *et al*., 2008; Yang and Fernando, 2021). ROS are indispensable for the activation of plant defense responses as a signaling component (Lee *et al*., 2020) or they can cause peroxidation of lipids, oxidation of proteins, damage to nucleic acids, enzyme inhibition, and activation of PCD pathway during hypersensitive response (Camagna and Takemoto, 2018), thus leading to the inhibition of biotrophic pathogens. However, the Fusarium infection represents another example, since ROS accumulation is induced by its toxins in the host plants (Desmond *et al*., 2008). Fusarium infection follows the H_2_O_2_ produced by the host cells, which causes enhanced production of fungal toxins enabling hyphae penetration into the host cells (Khaledi *et al*., 2016).

Secretory peroxidases were one of the most affected protein groups in both *HvMPK3* KO and WT lines after *F. graminearum* infection. All peroxidase isoforms were downregulated in WT, but three out of five peroxidases were upregulated in *HvMPK3* KO lines in response to *F. graminearum*. Secretory peroxidases belong to class III of plant peroxidases, having a broad spectrum of activities in plants (Almagro *et al*., 2009; Passardi *et al*., 2005; Sasaki *et al*., 2004). Such peroxidases belong to a PR protein 9 subfamily and are supposed to reduce the progression of pathogens by a generation of structural barriers. They are bifunctional because they exert either ROS-scavenging or ROS-producing activity (Passardi *et al*., 2004). Peroxidase activity contributes to the ROS generated during oxidative burst (O’Brien *et al*., 2012). ROS produced by peroxidases may either have signaling roles or contribute to the toxic environment for the pathogen (Camejo *et al*., 2016).

One of the crucial features of secretory peroxidases is to mediate cross-linking of cell wall components leading to cell wall reinforcement. They may cross-link extensins leading to the formation of large oligomers (Mishler-Elmore *et al*., 2021). The cross-linking of ferulic acid by covalent bonds requires the presence of cell wall peroxidases. Such cross-linked ferulic acid may create intra- or inter-polysaccharide bonds leading to cell wall stiffening (Fry, 2004). Lignin formation depends on oxidative cross-linking of monolignols (Ralph *et al*., 2004) and expression of peroxidases was shown to correlate with lignification (Cosio *et al*., 2017; Sorokan *et al*., 2014). Similarly, suberin poly-phenolic domain assembly requires peroxidase mediated oxidative coupling reactions (Bernards *et al*., 2004). Moreover, transient overexpression of *HvPrx40* a class III peroxidase in barley epidermis, improves resistance against *Blumeria graminis* f.sp. *hordei* pathogen (Johrde and Schweizer, 2008). It is likely that peroxidases similar to peroxidase 12 (*Arabidopsis thaliana*), cationic peroxidase PNC1 (*Arachis hypogaea*), and peroxidase HRPN (*Armoracia rusticana*) contribute to the formation of suberized structural barriers which exclude *F. graminearum* to the extracellular space in *HvMPK3* KO plants. Moreover, a genomic study reported a possible link of GDSL esterase/lipases to suberin biosynthesis (Soler *et al*., 2007). Protein similar to GDSL esterase/lipase At5g33370 (*Arabidopsis thaliana*) was significantly upregulated in *HvMPK3* KO plants. According to our results, *HvMPK3* KO lines activated a different set of LTPs compared to WT. A protein similar to lipid transfer-like protein VAS, implicated in suberin formation in Arabidopsis (Edstam *et al*., 2013), was upregulated in *HvMPK3* KO lines. Also, Pitzschke *et al*., (2014) found that AZI1 (belonging LTPs) directly interacts with AtMPK3, and confers salinity tolerance in Arabidopsis (Pitzschke *et al*., 2014). In summary, our results indicate that the cell wall stiffening by suberin in surface root cells might substantially contribute to the resistance of *HvMPK3* KO lines against *F. graminearum*. This might suggest a connection between HvMPK3 and LTPs as a part of barley defense mechanism against *F. graminearum* infection.

Since TALEN-based knockout of *HvMPK3* causes resistance to *F. graminearum* it provides a reliable tool for targeted biotechnological application in important cereal crop such as barley.

## Experimental procedures

### Cultivation of plant material

Three independent homozygous knock-out lines of barley (*Hordeum vulgare* L.) designated as *HvMPK3* KO-A, *HvMPK3* KO-B and *HvMPK3* KO-D, and control WT lines (Takáč *et al*., 2021) were used for the experiments. Seeds were surface sterilized using 5% (v/v) sodium hypochlorite and 70% (v/v) ethanol, followed by an additional incubation with 500 mM (v/v) H_2_O_2_ overnight. After sterilization, the seeds were imbibed and stratified on plates with agar-solidified nitrogen-free Fahräeus medium (Fåhraeus, 1957) at 4°C for synchronous germination (Perrine-Walker *et al*., 2007). The stratified sterile seeds on plates were transferred to phytotron (Weiss-Gallenkamp, Loughborough, United Kingdom) and cultivated at 21°C for 16 h in the light (day), and 8 h in darkness (night), 70% relative humidity with light levels of 150 μmol.m^-2^.s^-1^ provided by cool white fluorescent tubes (Philips Master tl-d 36W/840).

### Preparation of fluorescent F. graminearum

Mycelia of *F. graminearum* were cultivated in Mung bean soup (140 rpm, 20°C, 3 days), subsequently filtered through glass wool and centrifuged (300 rpm, room temperature, 5 min). Conidia obtained as such, were resuspended in sterile distilled water and inoculated on yeast extract-peptone-dextrose (YPD) medium. After overnight cultivation (180 rpm, 30°C), the mycelium was filtered through Miracloth (Merck) and washed with sterile distilled water, and 1.2 M KCl. For protoplast isolation, mycelia were cultivated in a solution containing 250 mg driselase, 1 mg chitinase, and 100 mg lysing enzyme from *Trichoderma harzianum* (all from Sigma-Aldrich). After 3 h (90 rpm, 30 °C), the enzyme-protoplast solution was run over the frit and the flow-through was centrifuged (3000 rpm, 4 °C, 10 min). Obtained pellet was washed with 1.2 M KCl, then resuspended in STC buffer (1.2 M sorbitol, 50 mM CaCl_2_).

Plasmid (5 μg of *pNDN-OGG* containing *GFP;* Schumacher, 2012) was mixed with 2x STC buffer, *F. graminearum* protoplasts, and 50% (w/v) polyethylene glycol (PEG) 4000. After 25 min on ice, 50% (w/v) PEG 4000 was added. Next, the mixture was incubated 10 min at RT, followed by addition of 1x STC buffer. Prepared samples were mixed with Regeneration agar containing nourseothricin (300 μg.ml^-1^, Jena BioScience, *pNDN-OGG*) or hygromycin (100 μg.ml^-1^, InvivoGen, *pNDH-OCT*). After 5 days at 20 °C, nourseothricin- and hygromycin-resistant transformants were transferred into complete medium (CM) agar plates containing antibiotics in the final concentration of 100 μg.ml^-1^.

Genomic DNA was extracted from lyophilized *F. graminearum* mycelia according to Cenis, (1992). Diagnostic PCR was performed using GoTaq G2 Flexi DNA polymerase according to a manufacturer’s protocol (Promega). Primers were designed using Primer3 0.4.0 software and synthesized by Sigma-Aldrich. The presence of *pNDN-OGG* or *pNDH-OCT* in *F. graminearum* transformants was checked with primers PoliC_fw (5’-CCCGGAAACTCAGTCTCCTT-3’) and TgluC_rev (5’-GTCTTCCGCTAAAACACCCC-3’) (1,295 bp) or PoliC_fw and Ttub_rev (5’-GAGGTGTGAGCATGGAAGTGATG-3’) (1,593 bp), respectively.

Homokaryots of *OE:GFP* transformants were obtained by single spore isolation. Briefly, fungi were cultivated in Mung bean soup (20°C, 140 rpm) and after 5 days, the cultures were filtered through glass wool and centrifuged (10 min, 3000 rpm, 4°C). Obtained conidia were resuspended in sterile distilled water and sprayed onto CM agar plates containing appropriate antibiotics in the final concentration of 100 μg/ml.

### Infection of barley plants with Fusarium

Fusarium grown in potato dextrose agar (PDA) medium for 2 weeks at 28 °C was washed with sterile distilled water containing 0.01% (v/v) Tween and filtered through sterile miracloth to obtain conidial spores (~1 × 10^5^ spores.ml^-1^) (Erayman *et al*., 2015). Five days old barley seedlings were infected by dipping in freshly prepared conidial suspension (sterile distilled water containing 0.01% (v/v) Tween was used as control) and put back to the media for incubation for further experiments. Microscopic, biochemical and proteomic analyses were performed on control and Fusarium-treated plants 24h and 48h post-infection.

### Phenotypic and microscopic analysis

For microscopic analysis, the roots of both WT and *HvMPK3* KO plants 24h and 48h after inoculation with GFP-labeled *F. graminearum* were used. Infected plants were grown in microscopic growth chambers (Nunc™ Lab-Tek™ II Chambered Coverglass (Thermofischer scientific, USA)) and observed using 488 nm excitation for GFP and the 517-527 nm emission using confocal laser scanning microscope LSM710 (Carl Zeiss, Germany). The images were color-coded for the analysis of the penetration of the infection hyphae in root epidermal cells using appropriate Zen Blue software function. The numbers of appressoria and penetration events were counted from the images and were compared among the lines by one-way ANOVA test at significance level at p < 0.05. Roots were stained with PI (1 mg.ml^-1^) for 10 min, washed with liquid Fahräeus medium for 1 min, and observed under spinning disk microscope (Carl Zeiss, Germany) equipped with Plan-Apochromat 20×/0.8 NA (Carl Zeiss, Germany) at 488 nm (for GFP) and 561 nm (for PI) with emission filters BP525/50 (for GFP) and BP629/62 (for PI). Image post-processing was done using ZEN 2014 software (Carl Zeiss, Germany), and the percentage of the dead cells were calculated from the processed images using the formulae (Number of dead cells / total number of cells in the field) × 100. To document root length, both treated and untreated plants were photographed by camera once per day for 10 days after the root inoculation with GFP-labeled *F. graminearum* spores. The experiment was performed in three biological replicates. Ten plants were used per replicate and treatment for each line. The statistical significance of treatment vs. control was deemed by one-way ANOVA test at significance level at p < 0.05.

### ROS staining

Control plants and treated plants from individual lines were used for ROS analysis 24 hrs postinoculation with GFP-labeled *F. graminearum* using the modified protocol of Kristiansen *et al*., (2009). Mock-treated and infected roots of WT and *HvMPK3* KO plants were incubated in 30 μM 2’,7’-dichlorodihydrofluorescein diacetate (CM-H_2_DCFDA; Cat no. C6827, Invitrogen™, USA) and PI (1 mg.ml^-1^) diluted in Fahräeus medium. Seedlings in microscopic chambers were stained by perfusion, followed by incubation in darkness for 15 min. After incubation the residual stain was washed using sterile Fahräeus medium for 3 times using perfusion. After washing, the signal excited at 488 nm for ROS/GFP and 543 nm for PI was recorded at the emission range 517-527 nm for ROS/GFP and 610-625 nm (For PI) using confocal laser scanning microscope LSM710 (Carl Zeiss, Germany). The imaging of ROS accumulation in whole root was performed after staining with H_2_DCFDA using epifluorescence microscope Zeiss Axio Imager M2 (Carl Zeiss, Germany) with settings for GFP excitation and emission. ROS levels in roots were analyzed semi-quantitatively from the images with ZEN software (Carl Zeiss, Germany).

### Histochemical detection of suberin and lignin

Root segments (first 25 mm from the root apex) of WT and *HvMPK3* KO plants 24 h after inoculation with *F. graminearum* were used for suberin and lignin histochemical detection. Suberin and lignin staining was carried out according to a previously published protocol (Sexauer *et al*., 2021; Ursache *et al*., 2018). Freehand cross-sections of fixed and cleared root segments were made in the root region between 15 and 25 mm from the root apex and stained with 0,01% (v/v) Fluorol Yellow 088 in absolute ethanol (stock solution: 1% (w/v) Fluorol Yellow in DMSO) for 30 min. For lignin detection root sections were stained with 0,2% (w/v) Basic Fuchsin in ClearSee solution for overnight. Stained cross-sections of roots were observed under confocal laser scanning microscope LSM710 (Carl Zeiss, Germany) using following settings for suberin: excitation: 488 nm, detection: 500-550 nm and for lignin: excitation: 561 nm, detection: 600-650. Measurement of fluorescence intensity and post-processing were done using ZEN 2014 software (Carl Zeiss, Germany). Fluorescence mean intensities were calculated from a minimum of three biological replicates.

### Western Blot Analysis

Samples were prepared and western blots performed as described before (Takáč *et al*., 2021). Polyvinylidene difluoride (PVDF) membranes were incubated with rabbit primary anti-L-ascorbate peroxidase antibody (#AS 08 368), diluted 1:2000, or with rabbit primary anti-catalase antibody (#AS 09 501), diluted 1:2000, or with rabbit primary anti-PR-1 antibody (#AS 10 687, all from Agrisera, Sweden), diluted 1:2000, all in 1% (w/v) BSA in TBS-T at 4°C overnight. Membranes were washed thoroughly and subsequently incubated at room temperature with corresponding HRP-conjugated secondary antibody (Thermo Fisher Scientific), diluted 1:5000 in 1% (w/v) BSA in TBS-T for 1.5 h. Following three washing steps, membranes were incubated with commercial Clarity Western ECL Substrate (BioRad, Hercules, CA, United States) and documented in a ChemiDoc MP imaging system (BioRad). Experiments were performed in three biological replicates, and the statistical significance was evaluated using Student’s t test (p< 0.05).

### Analysis of peroxidase activities and spectrophotometric measurement of ROS

Specific activities of peroxidases were analyzed using staining on native PAGE gels as published previously (Takáč *et al*., 2016). ROS levels were estimated in water extracts using xylenol orange assay as described in Takáč *et al*., (2014). The analyses were carried out in three biological replicates on control and Fusarium-treated roots of WT and *HvMPK3* KO seedlings 24h post-inoculation. The statistical significance was evaluated using Student’s t test (p< 0.05).

### Relative protein quantitation using nano-Liquid Chromatography-Tandem Mass Spectrometry Analysis (nLC-MSMS)

Barley WT, and *HvMPK3* KO lines A, B, and D were used for proteomic study. Root parts from four plants of each line in control and treatment variants were pooled into one sample. Each sample was analyzed in two biological replicates, and the final differential proteomes of WT and *HvMPK3* KO lines were obtained by comparing 6 biological replicates of Fusarium-treated lines vs. corresponding mock controls. Proteins were extracted using phenol extraction and methanol/ammonium acetate precipitation, as described previously (Takáč *et al*., 2017). Total of 50 μg of proteins dissolved in 50 μl of 6 M urea were subjected to in-solution trypsin digestion.

The peptides were then cleaned on C18 cartridges (Bond Elut C18; Agilent Technologies, Santa Clara, CA), dried using SpeedVac, and utilized for nLC-MS/MS (Takáč *et al*., 2017).

Two micrograms of protein tryptic digest were analyzed as published previously (Takáč *et al*., 2021), using the Ultimate 3000 nano-LC system and LTQ-Ortbitrap Velos mass spectrometer (both Thermo Fisher Scientific). All raw and results files were deposited to a publicly accessible database (see Data Availability Statement for details). Protein identification and label-free quantification was performed by the Proteome Discoverer 2.1 (Thermo Fisher Scientific), and an in-house script based on precursor ion intensities, as described before (Takáč *et al*., 2021). Statistically significant results were filtered with ANOVA p ≤ 0.05, applied to proteins exhibiting the fold change ≥ 1.5.

Proteins identified by single peptide were excluded from the results. Proteins present in all six replicates corresponding to the control proteome, and absent in all the treated replicates were considered unique for the control proteome, and *vice versa*.

The differential proteomes were evaluated by gene onthology (GO) annotation analysis and screening of protein domains using OmixBox Functional analysis module, as specified previously (Takáč *et al*., 2021).

## Acknowledgement

We would like to thank Lena Studt from Department of Applied Genetics and Cell Biology, University of Natural Resources and Life Sciences, Vienna for helping us with *F. graminearum* transformation. We also thank technicians Petra Trčková, Pavlína Floková, Katarína Takáčová and Monika Vadovičová for their expert technical help in all stages of the presented work.

## Supporting Information

**Figure S1.** Colonization of roots in barley plants of WT and *HvMPK3* KO lines by *Fusarium graminearum* mycelia 10 days after inoculation with spores.

**Figure S2.** Characterization of distribution and growth pattern of *F. graminearum* hyphae in regard of the root longitudinal axis. **(A-D)** Cytospectre graphs showing a distribution pattern of fluorescent hyphae on the root surface of WT **(A)**, *HvMPK3* KO-A (**B**), HvMPK3 KO-B (**C**) and HvMPK3 KO-D (**D**) lines. **(E)** Quantitative assessment of anisotropy characterizing angular distribution of growing hyphae. **(F)** Quantitative assessment of fluorescence distribution skewness of hyphae on the surface of measured root area. Data are represented as mean ± SD; n = 10 images/line, error bars represent standard deviations. Different lowercase letters above the error bars represent statistical significance according to one-way ANOVA and subsequent LSD test at *p* value < 0.05.

**Figure S3.** Comparison of root phenotypes in barely WT and *HvMPK3* KO lines between nontreated and *Fusarium graminearum*-treated plants 10 days after inoculation with spores.

**Figure S4.** Fluorescence visualization of *F. graminearum* hyphae distribution on root surface of barley wild type (WT) and *HvMPK3* KO lines 48h after inoculation with spores. **(A, B)** WT root. **(C, D)** Root of *HvMPK3* KO-A line. **(E, F)** Root of *HvMPK3* KO-B line. **(G, H)** Root of *HvMPK3* KO-D line. Orthogonal view depicting root surface cell layers **(A, C, E, G)** and maximum intensity projection of the root surface **(B, D, F, H)**. Appressoria are indicated by asterisk and infection hyphae penetrating root epidermal cells are indicated by white arrows. Scale bars: 20 μm **(A-H)**.

**Figure S5.** Concentration of hydrogen peroxide in roots of WT and *HvMPK3* KO lines 24h after inoculation with *Fusarium graminearum* spores as examined by Xylenol orange assay.

**Figure S6.** Comparison of gene onthology annotation according to biological processes, carried out in differential proteomes of WT and *HvMPK3* KO plants treated by *Fusarium graminearum* for 24h.

**Figure S7.** Comparison of gene onthology annotation according to cellular compartments, carried out in differential proteomes of WT and *HvMPK3* KO plants treated by *Fusarium graminearum* for 24h.

**Figure S8.** Full scan of the entire original gel whose fragments are presented in Figure 8A.

**Figure S9.** Full scan of the entire original blot **(A)** and loading control **(B)** whose fragments are presented in Figure 8C.

**Figure S10.** Full scan of the entire original blot **(A)** and loading control **(B)** whose fragments are presented in Figure 8E.

**Figure S11.** Full scan of the entire original blot **(A)** and loading control **(B)** whose fragments are presented in Figure 8G. The highlighted regions show sections presented in Figure 8G.

**Table S1.** Summary and quantification details of differentially regulated proteins found in roots of wild type plants 24h after the treatment with *Fusarium graminearum*

**Table S2.** Summary and quantification details of differentially regulated proteins found in roots of HvMPK3 KO plants 24h after the treatment with *Fusarium graminearum*

**Table S3.** Proteins predicted to be localized in extracellular space.

## Authors contributions

JB, PV and MO conducted phenotypic and microscopic documentation and analysis. PV, TT and TP performed proteomics analysis. PV and PM conducted immunoblot analysis, native electrophoresis and spectrophotometric measurements. OŠ performed suberin and lignin staining and related microscopic observations and evaluation. PK selected *HvMPK3* KO and WT barley lines. MK performed Fusarium transformation. JB, PV, MO, TT, GK and JŠ drafted the manuscript with input from all co-authors. JŠ conceived and supervised the project, provided infrastructure and secured funding.

## Data availability

The mass spectrometry proteomics data have been deposited to the ProteomeXchange Consortium via the PRIDE partner repository with the dataset identifier PXD029964”.

Reviewer account details: Username, Password: t4gfjKBY

